# Bismuth nanoparticles obtained by a facile synthesis method exhibit antimicrobial activity against *Staphylococcus aureus* and *Candida albicans*

**DOI:** 10.1101/2020.06.05.137109

**Authors:** Roberto Vazquez-Munoz, M. Josefina Arellano-Jimenez, Jose Lopez-Ribot

**Affiliations:** The University of Texas at San Antonio, San Antonio, TX, USA; The University of Texas at Dallas. Richardson, TX, 75080

**Keywords:** Nanoantibiotics, bismuth nanoparticles, BiNPs, Biofilms, synthesis, BAL, antimicrobial nanomaterials, nanoantibiotics

## Abstract

Bismuth compounds are known for their activity against multiple microorganisms; yet, the antibiotic properties of bismuth nanoparticles (BiNPs) remain poorly explored. The objective of this work is to further the research of BiNPs for nanomedicine, particularly as a disinfectant and for future treatments. Stable PVP-coated BiNPs were produced by a chemical reduction process, in less than 30 minutes, in a heated alkaline glycine solution, by the chelation and reduction of the bismuth (III) ion; resulting in the generation of small, spheroid particles with a crystalline organization. We assessed the antibacterial and antifungal activity of bismuth nanoparticles. PVP-BiNPs showed potent antibacterial activity against the pathogenic bacterium *Staphylococcus aureus* and antifungal activity against the opportunistic pathogenic yeast *Candida albicans*, both under planktonic and biofilm growing conditions. Our results indicate that BiNPs represent promising antimicrobial nanomaterials, and this facile synthetic method may allow for further investigation of their activity against a variety of pathogenic microorganisms.

## INTRODUCTION

Bismuth (symbol: Bi, Z=83, A=208.98) is a metallic element, non-toxic for humans (Lethal Intake >5-20 g/day/Kg, for years) [1,2]. Bismuth is insoluble in water but soluble in some organic solutions. The water-solubility and lipophilicity of bismuth can be enhanced when it is complexed with lipophilic molecules [3] and its biocompatibility is increased when it is chelated with hydroxyl or sulfhydryl groups. Bismuth compounds have been used in medicine for more than two centuries [4]. Bismuth subsalicylate is utilized to treat diarrhea-related ailments since the 1900s [5], and most recently, bismuth compounds have been used in computed tomography imaging and anti-cancer therapy [6,7]. In the U.S., around 30% of the bismuth compounds are used for cosmetic and pharmaceutical applications [4].

Several bismuth compounds display antibacterial and antifungal activity [8,9], and can outperform the inhibitory activity of conventional antibiotics [10]. When bismuth is thiolated, such as in the case of bismuth-thiols (BTs) complexes, its antimicrobial activity is further enhanced [11]. Bismuth-dimercaptopropanol (bismuth-BAL) and other BTs display anti-biofilm activity [12]. Additionally, the thiolation of bismuth complexes increases their stability [13]; however, BTs stability is still relatively low when compared to other antimicrobial agents [14].

The stability and antimicrobial activity of bismuth can be increased using nanotechnological approaches, due to their special structural characteristics. Phan *et al* suggested that nanomaterials may display greater stability than their precursors under some specific conditions [15]. Also, the capping agents provide stability to nanoparticles [16]. Stability is critical for controlling the BTs potential toxicity [17] and for extending their shelf-life. In the last two decades, metallic elements with known –or potential-antimicrobial properties (silver, copper, and titanium, among others) have been used for synthesizing antimicrobial nanomaterials (nanoantibiotics) [18].

Although some nanoantibiotics have been successfully transitioned to the market [19,20], some promising metallic elements, such as bismuth, remain under-researched, even though some early reports indicate that bismuth nanoparticles (BiNPs) display promising antimicrobial activity on bacteria, fungi, and protozoan [3,21]. According to Web of Science, 300 studies on BiNPs have been published in the last 20 years, yet only 12 are related to their antimicrobial activity (**fig. 1**). This scarceness on research may be due to the difficulties in synthesizing BiNPs and their low stability under culture conditions.

**Fig. 1.**
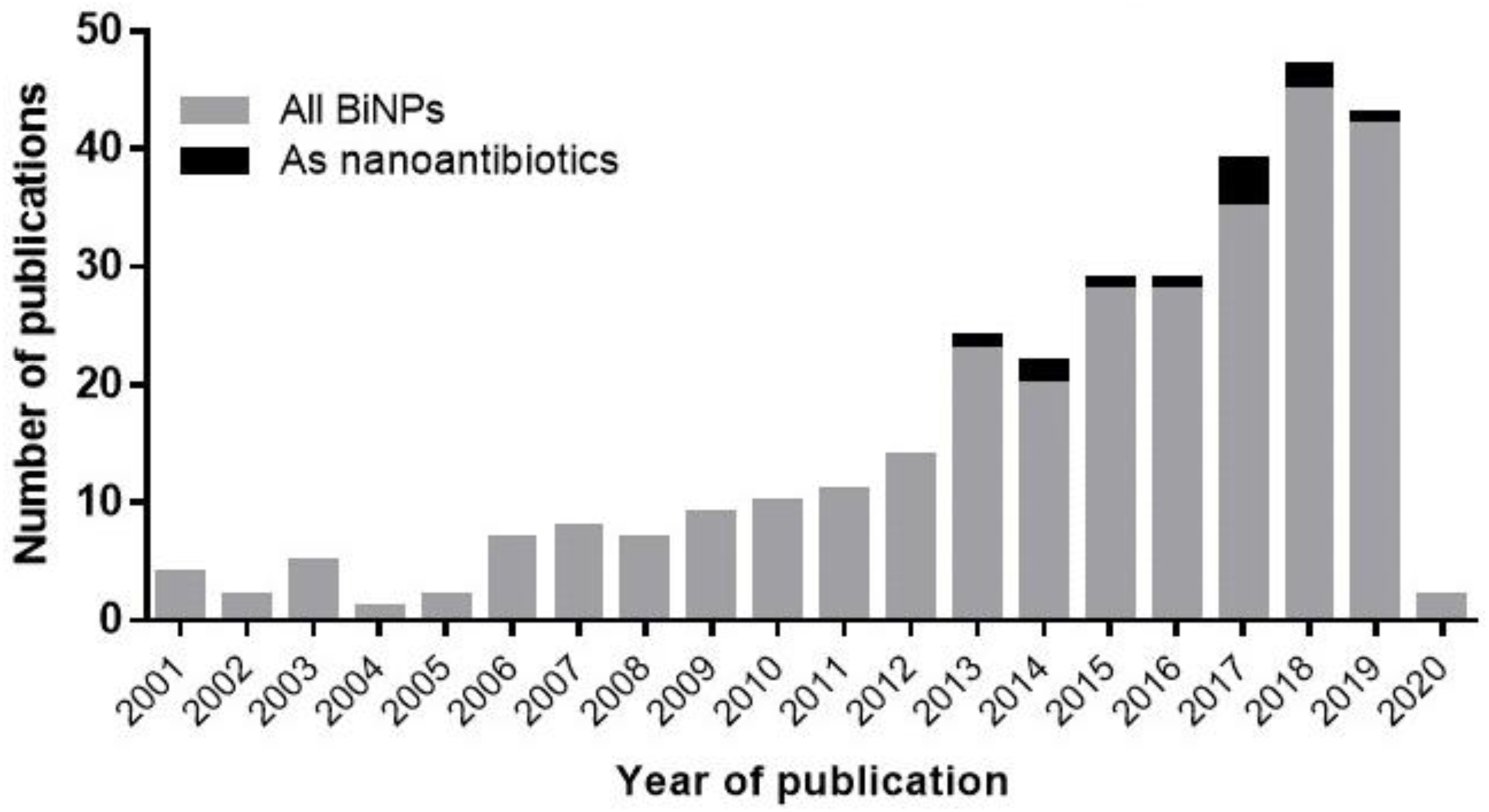
Research of BiNPs in recent literature. BiNPs as antimicrobial agents remain under-researched. According to Web of Science, approximately 300 studies on BiNPs have been published in the last 20 years, from which only 12 are related to their antimicrobial activity.

Most of the current protocols for synthesizing biologically suitable BiNPs cannot be replicated in non-specialized laboratories. Some methods require specialized equipment, such as gamma irradiators [22], laser ablation [23]; or controlled conditions, such as inert atmosphere [24], vacuum [25] or high temperatures [26,27]. Also, some protocols require a long-time synthesis [22], or several intermediate steps [28]. Although biosynthesis -green chemistry-has been explored, it is not easily replicable in non-specialized laboratories [29,30]. In general, BiNPs obtained using the above-mentioned protocols display an aspect ratio close to 1; although polyhedral, rod-like, and triangular shapes have also been described, and their sizes range from 1.7 nm to 1,000 nm (1 µm). Most recently our group has described a method for the fast, facile, and inexpensive synthesis of BiNPs using a simple chemical reduction process, which results in the generation of small nanoparticles with an aspect ratio close to one. This novel methodology does not require the use of sophisticated equipment, is easy-to-replicate, and can be easily implemented in non-specialized laboratories [31]. The main objective of this work was to characterize the antimicrobial activity of these easy-to-synthesize BiNPs against *Staphylococcus aureus*, a pathogenic bacterium, and *Candida albicans*, an opportunistic, dimorphic yeast; and in the process increasing the research on this new type of nanoantibiotics.

## MATERIALS AND METHODS

### Reagents and nanomaterials

2,3-Bis(2-methoxy-4-nitro-5-sulfophenyl)-2H-tetrazolium-5-carboxanilide salt (XTT), menadione, and phosphate Saline Buffer (PBS) from Sigma-Aldrich (MO). Osmium Tetroxide (OsO_4_) and Glutaraldehyde from Ted Pella; and PrestoBlue ™ Cell Viability Reagent from InVitrogen. 1% OsO_4_ and 2.5 % glutaraldehyde solutions were prepared in Milli Q water. A 10% PrestoBlue solution was prepared in BPS.

#### Nanoantibiotics

PVP-coated bismuth nanoparticles were produced by an easy-to-synthesize chemical reduction protocol recently reported by our group [31]. Briefly, in a pre-warmed glycine solution, bismuth nitrate salts were added, and the pH was raised to 9. Then, BAL and PVP solutions were consecutively added to the bismuth solution. Finally, a NaBH4 solution was added, dropwise, in two batches three minutes apart from each other. The suspension was kept on vigorous stirring for 10 minutes.

### Characterization of the PVP-BiNPs

In a previous report, we demonstrated the presence and essential characteristics of the BiNPs, via Transmission Electron Microscopy and Energy Dispersive X-ray Spectroscopy, as well as the ƺ-potential via Dynamic Light Scattering Spectroscopy [31]. In this study, we expanded the characterization of the BiNPs using different rounds of synthesis, as follows: *High-Resolution Transmission Electron Microscopy* (HR-TEM) for determining the single-particle structural lattice, using Selected-Area Electron Diffraction (SAED) analysis. The PVP-BiNPs, deposited on Type-B carbon-coated copper grids (Ted Pella Inc.), were examined in a JEOL 2010-F HR-TEM (Jeol Ltd.), with 200 kV accelerating voltage. *Dynamic Light Scattering analysis* to determine the hydrodynamic size, using a Zetasizer Nano ZS (Malvern Panalytical). PVP-BiNPs were diluted in Milli Q water and then transferred to a DTS1070 cell for the analysis. *UV-Vis spectroscopy* to follow the transition from bismuth (III) ions to the PVP-BiNPs, via the UV-Visible absorbance profile, in 10 nm steps, collected in a Genesys10 UV-Vis spectrophotometer (Thermo Fisher), from 200 to 600 nm.

### Antimicrobial susceptibility assays

#### Strains and culture conditions

In this study, the Gram-positive bacterium USA200 methicillin-sensitive *S. aureus* (MSSA) UAMS-1 strain and the dimorphic yeast *C. albicans* strain SC5314 were used. For growth, *S. aureus*, inoculated in tryptic soy broth (TSB) (BD Difco, MD) and *C. albicans*, inoculated in Yeast extract Peptone Dextrose (YPD) broth, were incubated overnight in an orbital shaker at 35°C.

#### BiNPs antimicrobial activity on the microbial planktonic cells

We followed the CLSI guidelines to evaluate the susceptibility to the BiNPs, with slight modifications. For *S. aureus* the CLSI M07 [32] guidelines were followed; whereas for *C. albicans* the CLSI M27 [33] guidelines were used. Briefly, overnight microbial cultures were washed twice in PBS; adjusted for a final concentration of 10^6^ cells mL^-1^ in MH broth for *S. aureus*; and to 10^3^ cells mL^-1^, in RPMI culture media, for *C. albicans*. Then, 50 µL of each microbial strain were transferred to 96-multiwell plates. Subsequently, bismuth compounds [BiNPs, bismuth-BAL and Bi(NO_3_)_3_] were prepared in a two-fold dilution series, for a final concentration range from 0.5 to 256 µg mL^-1^ (from 2.39 µM to 1.22 mM), and 50 µL were added to the multiwell plates with the microbial cells. Later, the multiwell plates were cultured at 180 rpm; at 37 °C for 24 h. The Minimal Inhibitory Concentration (MIC) was determined as the concentration at which no turbidity was detected.

#### Assessment of the BiNPs antibiofilm activity on S. aureus

The ability of the BiNPs to inhibit biofilm formation by *S. aureus* was evaluated following a protocol previously published by our group with minor modifications [34]. Briefly, a “*biofilm broth*” was prepared as follows: 45% Tryptic-Soy Broth (BD Difco, MD), 45 % Brain-Hearth Infusion broth (BD Difco, MD), and 10% Bovine Fetal Serum (BD Difco, MD). Bacterial cells were washed twice in PBS and adjusted to 10^8^ cells mL^-1^ in the *biofilm broth* and transferred to the 96-multiwell plates. Bismuth compounds were prepared in a two-fold dilution series, in the *biofilm broth*, for a final concentration range from 0.5 to 256 µg mL^-1^ (from 2.39 µM to 1.22 mM) and transferred to the multiwell plates with *S. aureus*. The plates were cultured at 37 °C for 24 h. After the incubation time, the biofilms were washed with PBS and the Presto Blue™ was added, and the multi-well plates were read spectrophotometrically as previously described by our group [35].

#### Assessment of the BiNPs antibiofilm activity on C. albicans

The anticandidal activity of BiNPs, for preventing the biofilm formation, was evaluated using the methods previously reported by our group [36], with minor modifications. *C. albicans* cells from overnight cultures were washed in PBS, adjusted to x10^6^ cells mL^-1^ in RPMI culture media, and transferred to the 96-multiwell plates. Then, BiNPs two-fold dilution series were prepared in RPMI, for a final concentration range from 0.5 to 256 µg mL^-1^, and transferred to the plates with *C. albicans*, and then cultured at 35 °C for 24 h. Post-incubation, *C. albicans* biofilms were washed with PBS, then XTT/menadione was added. The absorbance of XTT was collected in a Benchmark Microplate Reader (Bio-Rad Inc), at λ=490 nm.

To generate the dose-response curves, the normalized results from the absorbance readings (from *S. aureus* and *C. albicans*) were fit to the variable slope Hill equation (for the nonlinear drug dose-response relationship) using Prism 8 (GraphPad Software Inc). Then, the BiNPs IC_50_ was calculated. The IC_50_ was established as the concentration of BiNPs that reduce the 50 % of the bacterial growth. To ensure the reproducibility of the antimicrobial activity from the different rounds of BiNPs syntheses and the bismuth compounds, we used two biological replicates (multi-well plates) with two technical replicates within each plate.

### Ultrastructural analysis

To assess the impact of the BiNPs on the biofilm structure of *S. aureus* and *C. albicans*, the samples were analyzed via Scanning Electron Microscopy (SEM). Briefly, BiNPs-treated and control (untreated) samples were washed with PBS, fixed with a 2% glutaraldehyde solution for 60 minutes, and then stained and post-fixed with a 1% Osmium tetroxide solution, for 30 minutes at 4 °C. Next, samples were washed with PBS and dehydrated using an ascending concentration ethanol series, up to 100%, and left to dry overnight. Finally, the samples were coated with gold in a Sputter Coater SC7620 (Quorum Technologies), for 25 milliamperes for 3 minutes. The gold-coated samples were observed in a TM4000Plus Scanning Electron Microscope (Hitachi Inc.), with a magnification 500 and 2500x, using a 10 KeV voltage.

## RESULTS AND DISCUSSION

### Further characterization of the BAL-mediated PVP-BiNPs

#### High-Resolution Transmission Electron Microscopy (HR-TEM)

In a previous report, HR-TEM revealed the morphology of the BiNPs [31]. For this work, an extended characterization via HR-TEM was performed in other rounds of syntheses. The shape of the nanoparticles displayed an aspect ratio close to 1, although few specific shapes (i.e. polyhedrons) were observed (**fig. s1, supplementary materials**). The average diameter of the nanoparticles is 8.4 nm ±6.7 nm, ranging from 1.7 nm to 44.4 nm (n=1,159) (**fig. s1, inset, supplementary materials**). Approximately, 86.1% of the particle size was below 15 nm. Some larger individual nanoparticles (50-100 nm) and clusters of nanoparticles were also observed.

The single-particle HR-TEM analysis revealed the crystallinity of the PVP-BiNPs. Lattice distance measurements agreed with reported low-index planes for crystalline Bi with cubic, hexagonal, and orthorhombic cell. The crystalline lattice depicted in **figure 2A** shows distances of 0.289, 0.253, and 0.246 nm, which suggest a monoclinic cell (0.290 nm is the distance reported for the (2 1-1) plane in the orthorhombic cell) or misorientation. The SAED from the BAL-mediated PVP-BiNPs showed the appearance of a clear ring diffraction pattern, which confirmed the formation of crystalline bismuth nanostructures. Interestingly, the experimental d-spacing corroborates that BiNPs display a mixed arrangement, conformed by cubic and hexagonal phases. The d-spacing for the cubic lattice were 2.68, 1.91, and 1.54 Å (hkl 110, 200, and 211, respectively); whereas for the hexagonal lattice were 2.35, 1.65, and 1.44 Å (hkl 104, 024, and 122, respectively) (**Fig. 2B**), according to the data from the standard powder diffraction cards of the Joint Committee on Powder Diffraction Standards (JCPDS), bismuth files numbers #26-0214 (cubic phase), and #44-1246 (hexagonal phase).

**Fig. 2.**
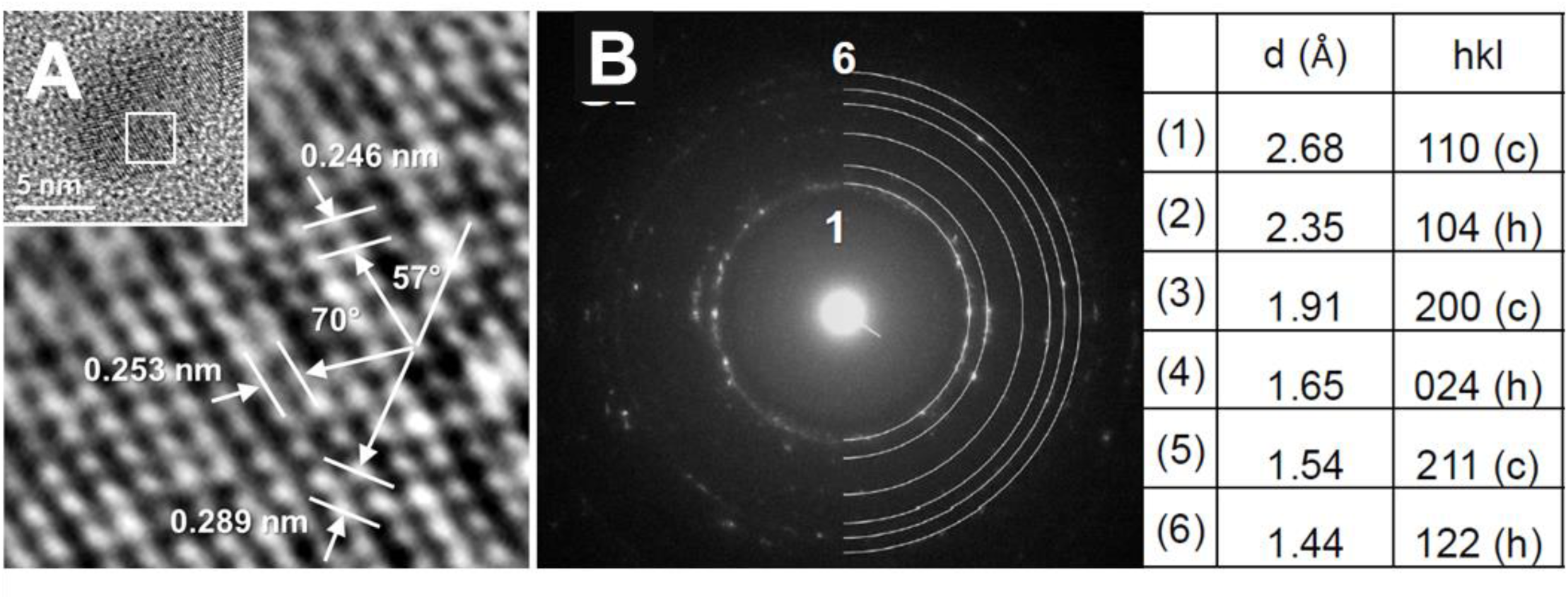
Further characterization of the BAL-mediated PVP-BiNPs. (A) HR-TEM analysis from a single particle confirms their crystalline organization, whereas the (B) Electron Diffraction Patter reveals their crystalline lattice as a cubic and hexagonal organization.

#### Dynamic Light Scattering spectroscopy

The PVP-BiNPs hydrodynamic size is 22.5 ±0.06 nm. The hydrodynamic size is greater than the metal core size observed on TEM images (22.5 nm vs 8.6 nm respectively). The increase in the hydrodynamic size may correspond to the 10K PVP coating chain-like molecules, whose structures extend when hydrated under the aqueous environment. The PVP coating improves BiNPs stability. HR-TEM images reveal that BiNPs are individual structures; however, they may be clustering over time due to the interactions of the organic compounds in the suspension.

#### UV-Vis characterization

The bismuth compounds displayed distinct absorbance profiles. The bismuth (III) ions profile (diluted in glycine) displayed an absorbance profile with a maximum at λ=240 nm, whereas the bismuth-BAL complex displayed a decreasing profile with a flat absorbance profile, from λ=340 to 390 nm, with a rapid decline λ=400 nm. In contrast, the BiNPs absorbance profile showed a surface plasmon with a shoulder at λ=250 nm, close to the reported by other authors [37], and a shoulder at λ=270 nm. Also, a flat profile was observed at λ=340 with a decrease at λ=380 nm. The difference observed in the absorbance profiles demonstrates the chemical transition from the bismuth (III) ions to the bismuth-BAL complex and then to the bismuth nanoparticles (**fig.3, bottom**). As seen in nanoparticles from other elements, the absorbance profile is determined by the surface physicochemical traits of the nanoparticles, such as shape and size distribution and the capping agents, among others [38]. In contrast to the AgNPs, a “typical” UV-Vis absorbance profile for BiNPs has still not been suggested. According to literature, different bismuth nanoparticles exhibit spectra profiles with maximum peaks at bands in different locations, such as λ=275 nm and λ=360 nm [39], λ=281 nm [40], 253 nm [37], and λ=313 nm [41]. Ma *et al* reported that BiNPs with a size below 10 nm did not display a UV-Vis absorbance profile. However, our small BiNPs did show a profile (average size <10 nm), yet, the absorbance profile may be from the BiNPs larger than 10 nm.

### Mechanism of BAL-mediated PVP-BiNPs synthesis

We used the bottom-up methodology approach for producing the BAL-mediated synthesis of the PVP-BiNPs. In a previous work, we described the procedure for synthesizing the nanoparticles [31]. Here, we propose the potential interactions between the precursors, aiming to describe the mechanism that leads to the formation of the PVP-BiNPs. The water-insoluble bismuth nitrate readily dissolves in glycine solution, allowing the release of bismuth (III) ions (**fig. 3A**), but the induction of an alkaline environment reduces the solubility of bismuth (III) ions. Yet, the alkaline environment allows dimercaptopropanol (BAL) to rapidly sequester the bismuth (III) ions, favoring the formation of the highly soluble bismuth-BAL complex (the solution changes from white to yellow, and the absorbance profile changes) (**fig. 3B**). PVP K-10 -a dispersant, mild reducing agent-, slowly initiates the formation of the nanoparticles, while regulates their size and shape. Moreover, PVP interacts chemically with bismuth [40], leading to the formation of a PVP coat on the nanoparticles. NaBH_4_ -a strong reducing agent-speed up the synthesis process. The rapid formation of the BiNPs is revealed by the color change of the reacting mixture (from yellow to black), and by a final transition on the UV-Vis absorbance (**fig. 3C**). The change of color to black is attributed to the increasing presence of –nanostructured-elemental bismuth, due to the shift of the oxidation state during the reduction of bismuth (III) ions [39]. The HR-TEM single-particle analysis and the DLS spectroscopy confirm the formation of bismuth nanoparticles, tentatively coated by PVP, as the hydrodynamic diameter is larger than the metallic BiNPs diameter.

**Fig. 3.**
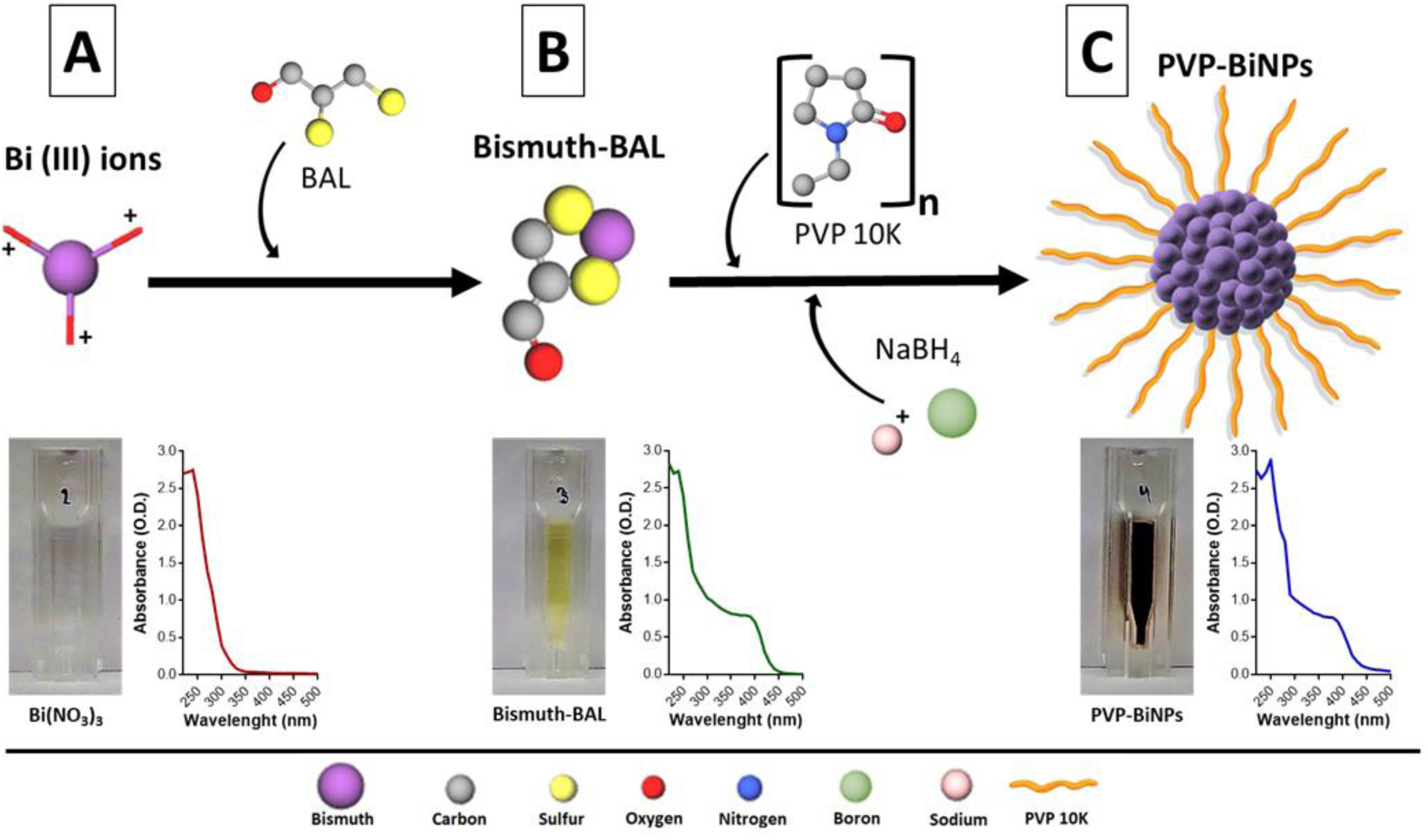
Proposed mechanism of synthesis for the formation of BiNPs. Bismuth (III) ions, solubilized in a glycine solution (A), interact with BAL, leading to the formation of the bismuth-BAL complex (B). Finally, NaBH_4_ induces the generation of PVP-BiNPs (C). Hydrogens atoms were not included in the molecular models for clarity. Molecules were built using the GLmol engine at http://molview.org/

### Stability of the PVP-coated BiNPs

The stability of bismuth nanoparticles in aqueous suspensions depends on several factors, such as pH [42], exposure to air [43], capping agent [44] and the bismuth-thiol bonding [11,13]. Thiolation increases the bismuth solubility and stability [13]; yet, the stability of bismuth-thiol complexes is still relatively low, as they remain in suspension for short periods [14]. Our synthesis method [31] enhances the BiNPs stability when compared with the bismuth (III) ions and the bismuth-BAL complex, under the same conditions of temperature, exposure to atmospheric air and changes on pH. Preparations of 15 mM bismuth nitrate and 15 mM bismuth-BAL complex precipitated within a few days; in contrast, the BAL-mediated PVP-coated BiNPs are stable for at least 11 weeks (**fig.s2, supplementary materials**). This increase in stability is an improvement over other processes for producing PVP-BiNPs, as it has been reported that PVP-coated BiNPs precipitate in less than 4 days [44]; Also, some studies show that BiNPs in aqueous suspensions are rapidly dissolved if the pH is out of the 8-10 range [42] or when exposed to air [45], yet our BiNPs remained stable for hours when dissolved in MilliQ water (pH 7) and when exposed to air. In our protocol, both solubility and stability of bismuth are increased due to the addition of BAL, leading to more stable BiNPs. The BAL-mediated synthesis of PVP-BiNPs kept the stability of nanoparticles even though the synthesis was performed under atmospheric air conditions.

Moreover, we tested variations on the synthesis method and found that PVP-BiNPs without BAL were unstable, precipitating in less than a week, whereas BiNPs without glycine or PVP were highly unstable, precipitating within minutes after the synthesis. These findings suggest that BiNPs stability is related to the bismuth (III) ions availability for the synthesis process (increased solubility) and capping molecules, with PVP acting as a capping/dispersant agent [46].

### BiNPs display antibacterial and antifungal activity on the planktonic and biofilms stages

PVP-BiNPs display strong antimicrobial activity against the planktonic cells *S. aureus* and *C. albicans* (**fig. 4A**). The BiNPs MIC against the planktonic cells of *S. aureus* was 1 µg mL^-1^, with a calculated IC_50_ of 0.28 µg mL^-1^. The antibacterial activity of these BiNPs is similar or better than the BiNPs activity reported before in the literature. Previously reported BiNPs MIC values ranged from 1.05 µg mL^-1^ for *S. mutans* and *S. gordonii* to 1,500 µg mL^-1^ for the MRSA strain [3,47]. Also, the antimicrobial activity of the resulting BiNPs is similar or better than several antimicrobial drugs, such as Vancomycin or Oxacillin (MIC= 1.5 µg mL^-1^) and Ceftaroline (MIC= 0.5 µg mL^-1^) [48]. Likewise, the BiNPs antimicrobial activity outperforms the antibacterial activity of silver nanoparticles (AgNPs) (MIC= 4 µg mL^-1^) [49].

**Fig. 4.**
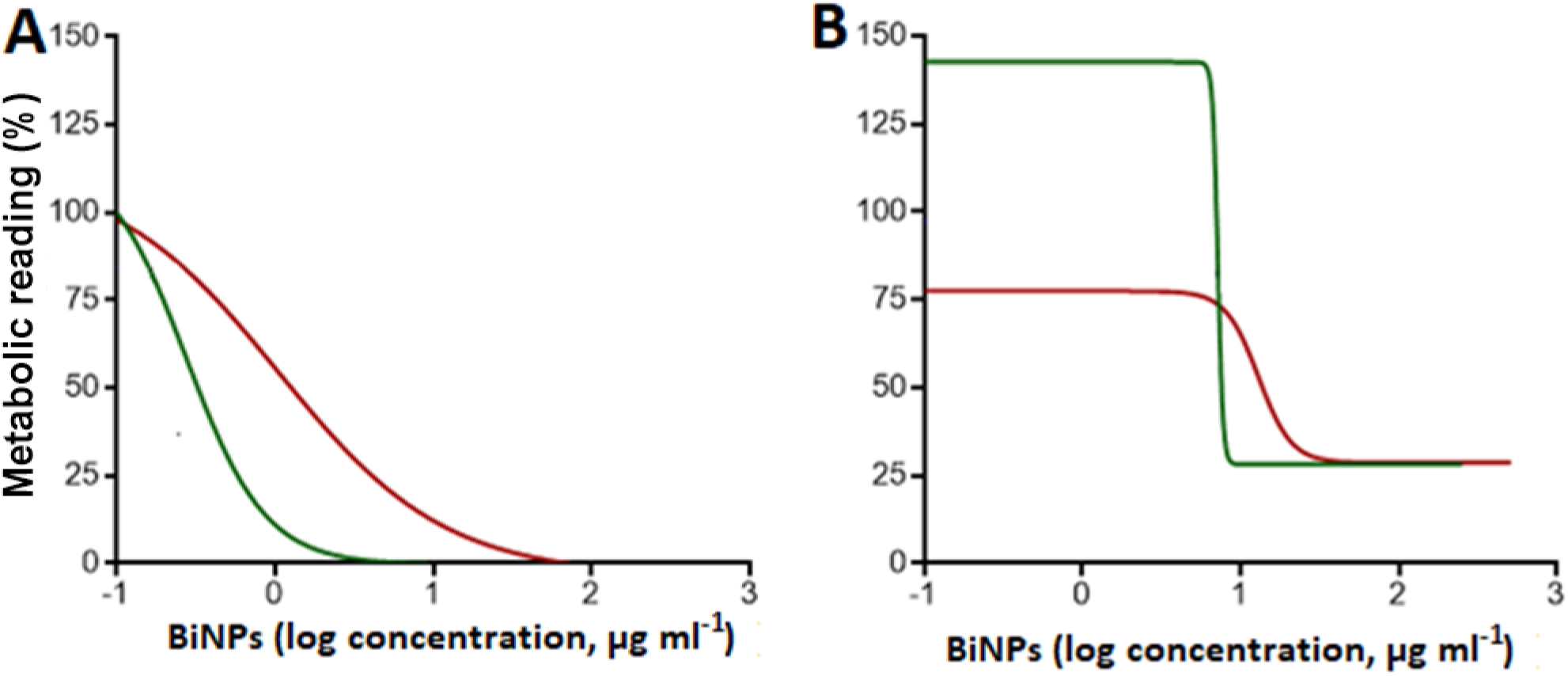
Antimicrobial activity of the BiNPs. Dose-response curves of the BiNPs antimicrobial activity on the planktonic cells (green line) and biofilms formation phase (red line) of the bacterium *S. aureus* (A) and the yeast *C. albicans* (B).

For *C. albicans*, the BiNPs MIC was 16 µg mL^-1^, whereas the calculated IC_50_ was 7.2 µg mL^-1^. As far as the authors know, the antifungal activity of BiNPs has only been addressed in one other study, also against *C. albicans*, with a MIC equivalent to 2.09 µg mL^-1^ [3]. When compared with antifungals, the BiNPs antimicrobial activity for *C. albicans* is parallel to azole antifungal agents but less potent than amphotericin B or echinocandins [50][51]. In contrast to their antibacterial activity, the BiNPs antifungal activity seems to be lower when compared with AgNPs (MIC= 2 µg mL^-1^) [49]

The synthesized PVP-BiNPs were also capable of inhibiting biofilm formation by both *S. aureus* and *C. albicans* (**fig. 4B**). In the case of *S. aureus* biofilms, the IC_50_ was 1.06 µg mL^-1^, which is among the most potent activity of BiNPs reported against the biofilm formation phase on bacteria. Baddidery *et al* reported that 2.61 µg mL^-1^ prevents the cell attachment on a polycarbonate membrane [52]. In contrast, Dalvand showed that even concentrations of BiNPs as high as 1,500 µg mL^-1^, inhibited the MRSA biofilms only by 16% [47]; Moreover, Hernandez-Delgadillo reported that a BiNPs concentration of 10,000 µg mL^-1^ was required to prevent the biofilm formation on *Enterococcus faecalis* [53]. When compared with antibacterial drugs, the BiNPs exhibit comparable antibiofilm activity to that of vancomycin (IC_100_=2 µg mL^-1^) and linezolid (IC_100_=4 µg mL^-1^) [54]

Regarding *C. albicans*, the calculated IC_50_ value for BiNPs against biofilms formed by this fungus was 7.9 µg mL^-1^, which interestingly is very close to its value under planktonic conditions As far as the authors know, the antibiofilm activity of BiNPs only has been addressed in one other study, where a concentration of 418 µg mL^-1^ was required to inhibit the *C. albicans* biofilm. [55]. When compared to antifungal agents. The activity of BiNPs is better than Fluconazole (IC_50_ >1024 µg mL^-1^), but lower than Amphotericin B (IC_50_ >1 µg mL^-1^) and the echinocandins [56,57].

Moreover, the antibacterial effect of Bi(NO_3_)_3_ and the bismuth-BAL complex was assessed against the planktonic stage and during the biofilm formation phase of *S. aureus* and *C. albicans*. For *S. aureus*, the antibacterial activity of the bismuth-BAL complex equals that of the BiNPs. In contrast, bismuth (III) ions have the lowest performance. For *C. albicans*, BiNPs displayed the highest antifungal activity, followed by the bismuth-BAL complex. Again, bismuth (III) ions exhibited the lowest activity. The activity of all bismuth compounds under planktonic and biofilm growing conditions is summarized in **table 1**, whereas the corresponding dose-response curves are in the supplementary materials section (**fig. s3**). As suggested by Domenico *et al* [58], the bismuth-BAL complex displays better antimicrobial activity than the Bi(NO_3_)_3_ salts. The antibacterial activity of different bismuth compounds against the bacterial planktonic stage and biofilm has been reported by different groups [8,12].

**Table 1.** Antimicrobial activity of bismuth compounds (µg mL^-1^) on the planktonic cells and biofilm stages of *S. aureus*.

| Phase/strain |  | Treatments Inhibitory concentration |  |  |
| --- | --- | --- | --- | --- |
|  |  | BiNPs | Bismuth-BAL complex | Bismuth (III) ions |
| Planktonic cells* | *S. aureus* | 1 | 1 | >256 |
|  | *C. albicans* | 16 | 64 | 128 |
| Biofilm formation** | *S. aureus* | 1.1 | 0.84 | 32 |
|  | *C. albicans* | 7.9 | >256 | >256 |

\* Minimal Inhibitory concentration (MIC)
\*\* Calculated $\text{IC}_{50}$

Furthermore, since the antimicrobial activity of BiNPs has been scarcely addressed, the mechanism of action has not been elucidated. Both BiNPs precursors –bismuth (III) ions and bismuth-BAL complex-display antimicrobial activity against both strains, therefore it is likely that the fundamental mechanisms of action of BiNPs are related to their precursors. The mechanisms of action of bismuth (III) ions are partially known. Ionic bismuth binds to glutathione, which improves its translocation inside of cells, from where it targets proteins and inhibit enzymes, such as urease and fumarase [59,60]. Regarding the bismuth-BAL complex, the mechanisms related to their antimicrobial activity are not fully understood. However, it is known that bismuth-thiols increase the permeability of the bacterial cell membrane [61]. Also, in the case of *S. epidermalis*, bismuth-thiol complexes reduce the glycocalyx production on the capsules [58]. It is highly likely that BiNPs release bismuth (III) ions with antimicrobial activity; though, due to the evident difference in the antimicrobial activity between BiNPs and bismuth (III) ions, the nanostructured arrangement of the BiNPs must be playing an unknown key role for enhancing the activity. So far, the mode of action of the BiNPs remains to be uncovered.

### The BAL-mediated PVP-BiNPs alter the biofilm ultrastructure

Scanning Electron Microscopy confirmed that BiNPs hinder the *S. aureus* ability to form biofilms. Low magnification SEM Images revealed that untreated *S. aureus* samples form biofilms with dense groups of cells (**fig. 5A**). High-magnification SEM images confirmed the presence of highly-packed multi-layered clusters of cells (**fig. 5D**). The morphology of the *S. aureus* cells revealed that they do not display any evident variations on their shape and size. In contrast, in the treated samples with sub-lethal concentrations of BiNPs (2 µg mL^-1^), the treated biofilms appeared to be less dense, but with a larger covered area (**fig. 5B**). The high-magnification images showed that the biofilm display lower thickness, with less clustered bacterial cells. Interestingly, the bacterial cell morphology was altered by the BiNPs, as evidenced by the diversity of cellular sizes and shapes observed (**fig. 5E**). This may be caused by the effects of bismuth on the cells, such as the denaturing of proteins [59,60] and changes in the cell membrane permeability [61]. Other potential mechanisms remain to be further studied and elucidated. In samples treated with higher concentrations of BiNPs (8 µg mL^-1^), the biofilms formation was almost completely abolished. Some sparse, small clusters of bacterial cells were present (**fig. 5C**), with most of the observed cells displaying similar shape and size, with the exception of a few larger cells (**fig. 5F**).

**Fig. 5.**
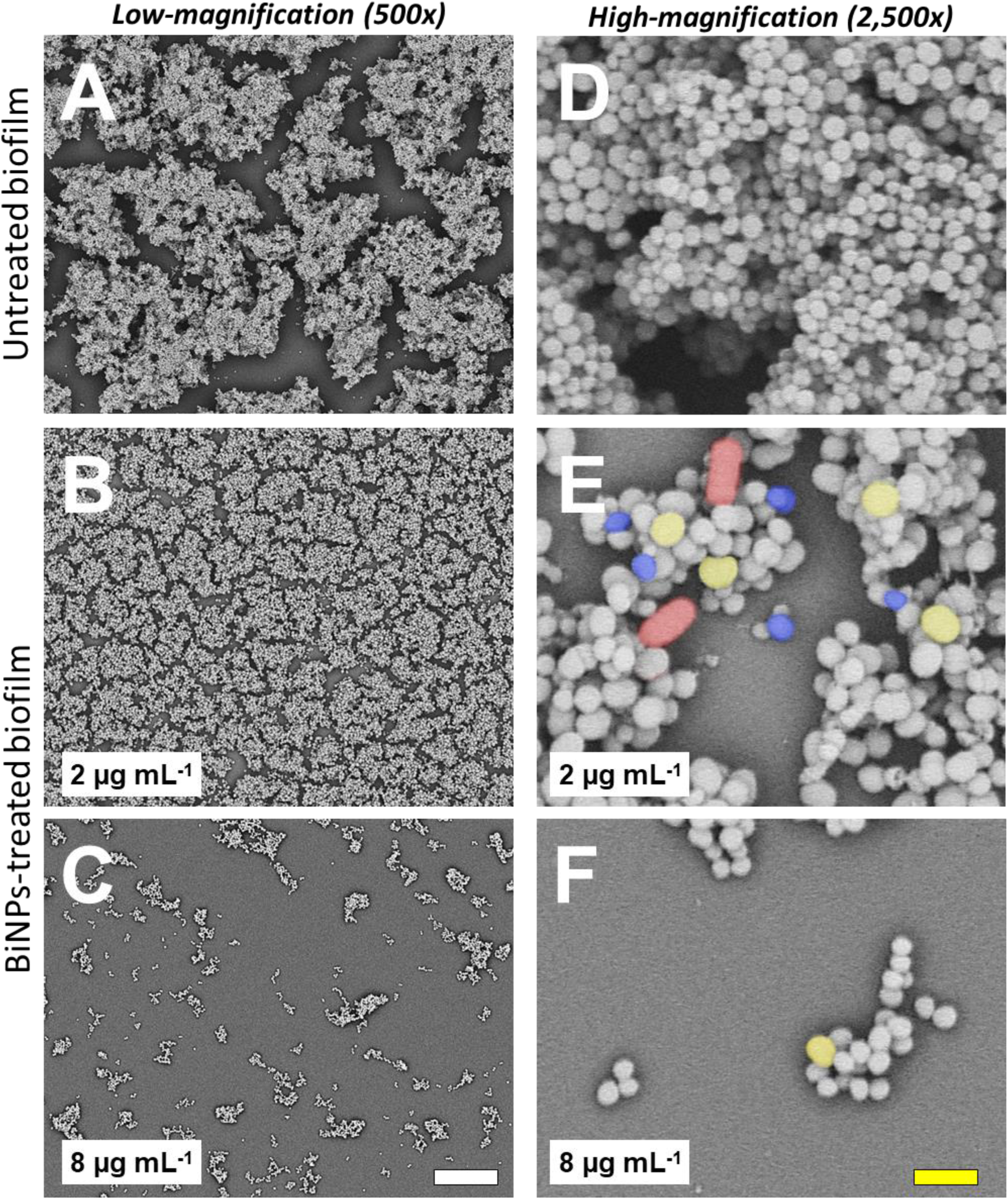
BiNPs inhibit biofilm formation by *S. aureus*. Untreated controls (A, D), as well as BiNPs-treated samples exposed to a low concentration of nanoparticles (B), form biofilms, but the cells from on the BiNPs-treated samples, display changes in morphology (E), from changes in size (yellow and blue-colored cells), and in shape (red-colored cells). Samples treated with a higher concentration of BiNPs do not form biofilms, but the cell morphology changes were less common (F). Scale bars: white=20 µm, yellow=2 µm.

Regarding the effect on *C. albicans* biofilms, SEM analysis confirmed that BiNPs negatively impact the dimorphic transition, from yeast to hyphae, and its ability to establish biofilms. On the untreated samples, low-magnification images revealed thick biofilms with fully-formed hyphae that completely cover the surface (**fig. 6A**). High-magnification images showed that only the hyphal shape is visible in the samples (**fig. 6D**). Samples treated with sub-lethal concentrations of BiNPs (4 µg mL^-1^), also showed thick biofilms that totally covered the surface (**fig. 6B**); however, alterations on the cell structure were evident. High-magnification images showed that yeast-like and pseudohyphae-like structures were abundant, confirming that BiNPs hinder the dimorphic transition ability of *C. albicans* (**fig. 6E**). The impact of bismuth nanoparticles on the ability of *C. albicans* to form biofilms became more evident in samples treated with higher concentrations of BiNPs (64 µg mL^-1^). At this concentration, biofilm formation was greatly inhibited by the BiNPs, as seen in **fig. 6C**, where only a few, aberrant-shaped hyphae were observable. The yeast-like and pseudohyphae-like structures also presented alterations on their morphology (**fig. 6F**).

**Fig. 6.**
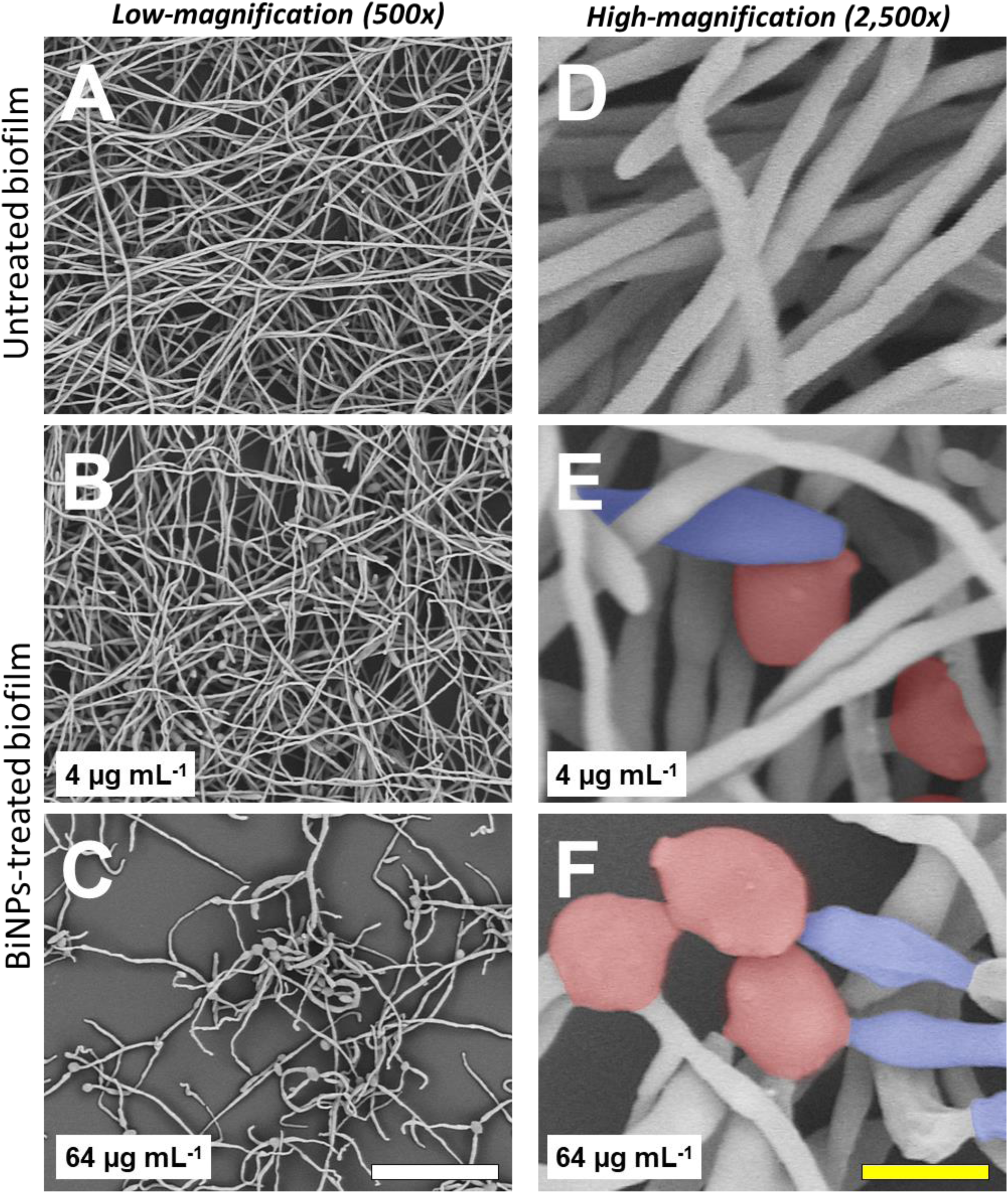
BiNPs affect the dimorphic transition and inhibit biofilm formation in *C. albicans*. Untreated controls (A, D) show the typical filamentous *C. albicans* biofilms, however, in samples treated with a low concentration of BiNPs, the dimorphic transition is affected. (B, E), In samples treated with a high concentration of BiNPs, biofilm formation is mostly inhibited, and the cells show aberrant morphologies (C, F). Yeast-like (in red) and pseudohyphae-like (in blue) cells were observed. Scale bars: white=40 µm, yellow=4 µm.

## CONCLUSION

Our small, spheroid PVP-coated bismuth nanoparticles are more stable than the bismuth-BAL complex in aqueous solutions. Also, these BiNPs display antimicrobial activity against the bacterium *S. aureus* and the yeast *C. albicans* under both planktonic and biofilm formation stages. The antimicrobial activity of these BiNPs is comparable to or better than other BiNPs synthesized by more sophisticated methods. Moreover, the antimicrobial activity of the BiNPs is parallel to the commercially available antibiotics and is also similar in potency to that of silver nanoparticles. Microscopy analysis reveals that BiNPs reduce biofilm formation and negatively alter the cell morphology in both *S. aureus* and *C. albicans*. This work shows that cost-effective, fast, easy-to-synthesize nanomaterials may display broad antimicrobial activity against bacteria and fungi. These nanomaterials could be applied as broad-range sanitizers to decrease the dispersion of potential pathogenic cells and for reducing their ability to form biofilms in healthcare-related facilities and other public spaces.

## ACKNOWLEDGMENTS

RV-M acknowledges the receipt of a postdoctoral scholarship from the CONACYT and the UTSA. Support in the laboratory was provided by the Margaret Batts Tobin Foundation, San Antonio, TX, USA (to JLL-R). the authors also thank the Graphic Designer Salma Carballo, for her assistance with the images.

## CONFLICT OF INTEREST

The authors declare there is no conflict of interest. The funders had no role in study design, data collection, data analysis, decision to publish, or preparation of the manuscript

## Notes

### Competing Interest Statement

The authors have declared no competing interest.

## REFERENCES

1. Ma L, Wu J, Wang S, Yang H, Liang D, Lu Z. Synergistic antibacterial effect of Bi2S3nanospheres combined with ineffective antibiotic gentamicin against methicillin-resistant Staphylococcus aureus. J Inorg Biochem. 2017;168: 38–45. doi: 10.1016/j.jinorgbio.2016.12.005

2. Sun H. Biological Chemistry of Arsenic, Antimony and Bismuth. Biological Chemistry of Arsenic, Antimony and Bismuth. Wiley; 2010. doi: 10.1002/9780470975503

3. Badireddy AR, Hernandez-Delgadillo R, Sánchez-Nájera RI, Chellam S, Cabral-Romero C. Synthesis and characterization of lipophilic bismuth dimercaptopropanol nanoparticles and their effects on oral microorganisms growth and biofilm formation. J Nanoparticle Res. 2014;16: 2456. doi: 10.1007/s11051-014-2456-5

4. USGS. Bismuth Statistics and Information. In: National Minerals Information Center [Internet]. 2019 [cited 14 May 2019]. Available: https://www.usgs.gov/centers/nmic/bismuth-statistics-and-information

5. Goldman RD. Bismuth salicylate for diarrhea in children. Can Fam Physician. 2013;59: 843–4. Available: http://www.ncbi.nlm.nih.gov/pubmed/23946025

6. Kowalik M, Masternak J, Barszcz B. Recent Research Trends on Bismuth Compounds in Cancer Chemoand Radiotherapy. Curr Med Chem. 2019;26: 729–759. doi: 10.2174/0929867324666171003113540

7. Cheng Y, Chang Y, Feng Y, Jian H, Tang Z, Zhang H. Deep-Level Defect Enhanced Photothermal Performance of Bismuth Sulfide-Gold Heterojunction Nanorods for Photothermal Therapy of Cancer Guided by Computed Tomography Imaging. Angew Chemie Int Ed. 2018;57: 246–251. doi: 10.1002/anie.201710399

8. Mahony DE, Lim-Morrison S, Bryden L, Faulkner G, Hoffman PS, Agocs L, et al. Antimicrobial activities of synthetic bismuth compounds against Clostridium difficile. Antimicrob Agents Chemother. 1999;43: 582–588. doi: 10.1128/aac.45.5.1417-1421.2001

9. Ferraz KSO, Reis DC, Da Silva JG, Souza-Fagundes EM, Baran EJ, Beraldo H. Investigation on the bioactivities of clioquinol and its bismuth(III) and platinum(II, IV) complexes. Polyhedron. 2013;63: 28–35. doi: 10.1016/j.poly.2013.07.008

10. Folsom JP, Baker B, Stewart PS. In vitro efficacy of bismuth thiols against biofilms formed by bacteria isolated from human chronic wounds. J Appl Microbiol. 2011;111: 989–996. doi: 10.1111/j.1365-2672.2011.05110.x

11. Domenico P, Salo RJ, Novick SG, Schoch PE, Van Horn K, Cunha BA. Enhancement of bismuth antibacterial activity with lipophilic thiol chelators. Antimicrob Agents Chemother. 1997;41: 1697–1703. doi: 10.1128/aac.41.8.1697

12. Varposhti M, Abdi Ali A, Mohammadi P. Synergistic effects of bismuth thiols and various antibiotics against Pseudomonas aeruginosa Biofilm. Jundishapur J Microbiol. 2014;7. doi: 10.5812/jjm.9142

13. Warren SC, Jackson AC, Cater-Cyker ZD, Disalvo FJ, Wiesner U. Nanoparticle Synthesis via the Photochemical Polythiol Process Scheme 1. Synthetic Route for the Photochemical Polythiol Process (Metals Tested Include Bismuth, Copper, Antimony, and Lead). J AM CHEM SOC. 2007;129: 35. doi: 10.1021/ja0733639

14. Rudramurthy GR, Swamy MK, Sinniah UR, Ghasemzadeh A. Nanoparticles: Alternatives against drug-resistant pathogenic microbes. Molecules. 2016;21: 1–30. doi: 10.3390/molecules21070836

15. Phan HT, Haes AJ. What Does Nanoparticle Stability Mean? 2019 [cited 30 Mar 2020]. doi: 10.1021/acs.jpcc.9b00913

16. Tejamaya M, Merrifield RC, Lead JR. Stability of Citrate, PVP, and PEG Coated Silver Nanoparticles in Ecotoxicology Media. 2012; 7011–7017. doi: 10.1021/es2038596

17. Auffan M, Achouak W, Rose J, Roncato MA, Chanéac C, Waite DT, et al. Relation between the redox state of iron-based nanoparticles and their cytotoxicity toward Escherichia coli. Environ Sci Technol. 2008;42: 6730–6735. doi: 10.1021/es800086f

18. Edson JA, Kwon YJ. Design, challenge, and promise of stimuli-responsive nanoantibiotics. Nano Converg. 2016;3: 26. doi: 10.1186/s40580-016-0085-7

19. Vance ME, Kuiken T, Vejerano EP, McGinnis SP, Hochella MF, Hull DR. Nanotechnology in the real world: Redeveloping the nanomaterial consumer products inventory. Beilstein J Nanotechnol. 2015;6: 1769–1780. doi: 10.3762/bjnano.6.181

20. Weissig V, Pettinger TK, Murdock N. Nanopharmaceuticals (part 1): products on the market. International journal of nanomedicine. 2014. doi: 10.2147/IJN.S46900

21. Vega-Jiménez AL, Almaguer-Flores A, Flores-Castañeda M, Camps E, Uribe-Ramírez M, Aztatzi-Aguilar OG, et al. Bismuth subsalicylate nanoparticles with anaerobic antibacterial activity for dental applications. Nanotechnology. 2017;28: 435101. doi: 10.1088/1361-6528/aa8838

22. El-Batal AI, El-Sayyad GS, El-Ghamry A, Agaypi KM, Elsayed MA, Gobara M. Melanin-gamma rays assistants for bismuth oxide nanoparticles synthesis at room temperature for enhancing antimicrobial, and photocatalytic activity. J Photochem Photobiol B Biol. 2017;173: 120–139. doi: 10.1016/j.jphotobiol.2017.05.030

23. Reus TL, Machado TN, Bezerra AG, Marcon BH, Paschoal ACC, Kuligovski C, et al. Dose-dependent cytotoxicity of bismuth nanoparticles produced by LASiS in a reference mammalian cell line BALB/c 3T3. Toxicol Vitr. 2018;53: 99–106. doi: 10.1016/J.TIV.2018.07.003

24. Gomez C, Hallot G, Pastor A, Laurent S, Brun E, Sicard-Roselli C, et al. Metallic bismuth nanoparticles: Towards a robust, productive and ultrasound assisted synthesis from batch to flow-continuous chemistry. Ultrason Sonochem. 2019;56: 167–173. doi: 10.1016/j.ultsonch.2019.04.012

25. Bi H, He F, Dong Y, Yang D, Dai Y, Xu L, et al. Bismuth Nanoparticles with “Light” Property Served as a Multifunctional Probe for X-ray Computed Tomography and Fluorescence Imaging. Chem Mater. 2018;30: 3301–3307. doi: 10.1021/acs.chemmater.8b00565

26. Winter H, Christopher-Allison E, Brown AL, Goforth AM. Aerobic method for the synthesis of nearly size-monodisperse bismuth nanoparticles from a redox non-innocent precursor. Nanotechnology. 2018;29: 155603. doi: 10.1088/1361-6528/aaacb9

27. Wei B, Zhang X, Zhang C, Jiang Y, Fu YY, Yu C, et al. Facile Synthesis of Uniform-Sized Bismuth Nanoparticles for CT Visualization of Gastrointestinal Tract in Vivo. ACS Appl Mater Interfaces. 2016;8: 12720–12726. doi: 10.1021/acsami.6b03640

28. Reverberi A Pietro, Varbanov PS, Lauciello S, Salerno M, Fabiano B. An eco-friendly process for zerovalent bismuth nanoparticles synthesis. J Clean Prod. 2018;198: 37–45. doi: 10.1016/J.JCLEPRO.2018.07.011

29. Shakibaie M, Amiri-Moghadam P, Ghazanfari M, Adeli-Sardou M, Jafari M, Forootanfar H. Cytotoxic and antioxidant activity of the biogenic bismuth nanoparticles produced by Delftia sp. SFG. Mater Res Bull. 2018;104: 155–163. doi: 10.1016/J.MATERRESBULL.2018.04.001

30. Nazari P, Faramarzi MA, Sepehrizadeh Z, Mofid MR, Bazaz RD, Shahverdi AR. Biosynthesis of bismuth nanoparticles using Serratia marcescens isolated from the Caspian Sea and their characterisation. IET Nanobiotechnology. 2012;6: 58. doi: 10.1049/iet-nbt.2010.0043

31. Vazquez-Munoz R, Arellano-Jimenez MJ, Lopez-Ribot JL. Fast, facile synthesis method for BAL-mediated PVP-bismuth nanoparticles. MethodsX. 2020;7: 100894. doi: 10.1016/j.mex.2020.100894

32. CLSI. M07. Methods for Dilution Antimicrobial Susceptibility Tests for Bacteria That Grow Aerobically. 11th ed. Weinstein MP, editor. Wayne, PA; 2018. Available: https://clsi.org/standards/products/microbiology/documents/m07/

33. CLSI. M27. Reference Method for Broth Dilution Antifungal Susceptibility Testing of Yeasts. 4th ed. Alexander BD, editor. Wayne, PA: Clinical Laboratory Standards Institute; 2017. Available: https://clsi.org/standards/products/microbiology/documents/m27/

34. Torres NS, Abercrombie JJ, Srinivasan A, Lopez-Ribot JL, Ramasubramanian AK, Leung KP. Screening a Commercial Library of Pharmacologically Active Small Molecules against Staphylococcus aureus Biofilms. Antimicrob Agents Chemother. 2016;60: 5663–5672. doi: 10.1128/AAC.00377-16

35. Montelongo-Jauregui D, Srinivasan A, Ramasubramanian AK, Lopez-Ribot JL. An in vitro model for oral mixed biofilms of Candida albicans and Streptococcus gordonii in synthetic saliva. Front Microbiol. 2016;7. doi: 10.3389/fmicb.2016.00686

36. Pierce CG, Uppuluri P, Tristan AR, Wormley FL, Mowat E, Ramage G, et al. A simple and reproducible 96-well plate-based method for the formation of fungal biofilms and its application to antifungal susceptibility testing. Nat Protoc. 2008;3: 1494–1500. doi: 10.1038/nprot.2008.141

37. Gomez C, Hallot G, Port M. Bismuth metallic nanoparticles. Inorganic Frameworks as Smart Nanomedicines. Elsevier; 2018. pp. 449–487. doi: 10.1016/B978-0-12-813661-4.00010-9

38. Baset S, Akbari H, Zeynali H, Shafie M. Size measurement of metal and semiconductor nanoparticles via UV-Vis absorption spectra. Dig J Nanomater Biostructures. 2011;6: 709–716.

39. Li J, Fan H, Chen J, Liu L. Synthesis and characterization of poly(vinyl pyrrolidone)-capped bismuth nanospheres. Colloids Surfaces A Physicochem Eng Asp. 2009;340: 66–69. doi: 10.1016/j.colsurfa.2009.03.007

40. Wang YW, Hong H, Kim KS. Size Control of Semimetal Bismuth Nanoparticles and the UV-Visible and IR Absorption Spectra. 2005 [cited 29 May 2019]. doi: 10.1021/jp046423v

41. Ma D, Zhao J, Li Y, Su X, Hou S, Zhao Y, et al. Organic molecule directed synthesis of bismuth nanostructures with varied shapes in aqueous solution and their optical characterization. Colloids Surfaces A Physicochem Eng Asp. 2010;368: 105–111. doi: 10.1016/j.colsurfa.2010.07.022

42. Brown AL, Goforth AM. PH-dependent synthesis and stability of aqueous, elemental bismuth glyconanoparticle colloids: Potentially biocompatible X-ray contrast agents. Chem Mater. 2012;24: 1599–1605. doi: 10.1021/cm300083j

43. Wang F, Tang R, Yu H, Gibbons PC, Buhro WE. Size- and Shape-Controlled Synthesis of Bismuth Nanoparticles. Chem Mater. 2008;20: 3656–3662. doi: 10.1021/cm8004425

44. Borovikova LN, Polyakova I V., Korotkikh EM, Lavrent’ev VK, Kipper AI, Pisarev OA. Synthesis and Stabilization of Bismuth Nanoparticles in Aqueous Solutions. Russ J Phys Chem A. 2018;92: 2253–2256. doi: 10.1134/S0036024418110055

45. Luz A, Feldmann C. Reversible photochromic effect and electrochemical voltage driven by light-induced Bi0-formation. J Mater Chem. 2009;19: 8107. doi: 10.1039/b907146f

46. Koczkur KM, Mourdikoudis S, Polavarapu L, Skrabalak SE. Polyvinylpyrrolidone (PVP) in nanoparticle synthesis. Dalt Trans. 2015;44: 17883–17905. doi: 10.1039/C5DT02964C

47. Firouzi Dalvand L, Hosseini F, Moradi Dehaghi S, Siasi Torbati E. Inhibitory Effect of Bismuth Oxide Nanoparticles Produced by Bacillus licheniformis on Methicillin-Resistant Staphylococcus aureus Strains (MRSA). Iran J Biotechnol. 2018;16: 279–286. doi: 10.21859/ijb.2102

48. Meeker DG, Beenken KE, Mills WB, Loughran AJ, Spencer HJ, Lynn WB, et al. Evaluation of Antibiotics Active against Methicillin-Resistant Staphylococcus aureus Based on Activity in an Established Biofilm. 2016 [cited 24 Feb 2020]. doi: 10.1128/AAC.01251-16

49. Vazquez-Muñoz R, Arellano-Jimenez MJ, Lopez FD, Lopez-Ribot JL. Protocol optimization for a fast, simple and economical chemical reduction synthesis of antimicrobial silver nanoparticles in non-specialized facilities. BMC Res Notes. 2019;12: 773. doi: 10.1186/s13104-019-4813-z

50. Shrestha SK, Fosso MY, Garneau-Tsodikova S. A combination approach to treating fungal infections. Sci Rep. 2015;5. doi: 10.1038/srep17070

51. Rautemaa R, Richardson M, Pfaller MA, Perheentupa J, Saxén H. Activity of amphotericin B, anidulafungin, caspofungin, micafungin, posaconazole, and voriconazole against Candida albicans with decreased susceptibility to fluconazole from APECED patients on long-term azole treatment of chronic mucocutaneous candidiasis. Diagn Microbiol Infect Dis. 2008;62: 182–185. doi: 10.1016/j.diagmicrobio.2008.05.007

52. Badireddy AR, Marinakos SM, Chellam S, Wiesner MR. Lipophilic nano-bismuth inhibits bacterial growth, attachment, and biofilm formation. Surf Innov. 2013;1: 181–189. doi: 10.1680/si.13.00009

53. Hernandez-Delgadillo R, Del Angel-Mosqueda C, Solís-Soto JM, Munguia-Moreno S, Pineda-Aguilar N, Sánchez-Nájera RI, et al. Antimicrobial and antibiofilm activities of MTA supplemented with bismuth lipophilic nanoparticles. Dent Mater J. 2017;36: 503–510. doi: 10.4012/dmj.2016-259

54. Cirioni O, Giacometti A, Ghiselli R, Bergnach C, Orlando F, Mocchegiani F, et al. Pre-treatment of central venous catheters with the cathelicidin BMAP-28 enhances the efficacy of antistaphylococcal agents in the treatment of experimental catheter-related infection. Peptides. 2006;27: 2104–2110. doi: 10.1016/j.peptides.2006.03.007

55. Hernandez-Delgadillo R, Velasco-Arias D, Martinez-Sanmiguel JJ, Diaz D, Zumeta-Dube I, Arevalo-Niño K, et al. Bismuth oxide aqueous colloidal nanoparticles inhibit Candida albicans growth and biofilm formation. Int J Nanomedicine. 2013;8: 1645–1652. doi: 10.2147/IJN.S38708

56. Nadhe SB, Singh R, Wadhwani SA, Chopade BA. Acinetobacter sp. mediated synthesis of AgNPs, its optimization, characterization and synergistic antifungal activity against C. albicans. J Appl Microbiol. 2019;127: 445–458. doi: 10.1111/jam.14305

57. Nett JE, Crawford K, Marchillo K, Andes DR. Role of Fks1p and matrix glucan in Candida albicans biofilm resistance to an echinocandin, pyrimidine, and polyene. Antimicrob Agents Chemother. 2010;54: 3505–3508. doi: 10.1128/AAC.00227-10

58. Domenico P, Baldassarri L, Schoch PE, Kaehler K, Sasatsu M, Cunha BA. Activities of bismuth thiols against staphylococci and staphylococcal biofilms. Antimicrob Agents Chemother. 2001;45: 1417–1421. doi: 10.1128/AAC.45.5.1417-1421.2001

59. Sadler PJ, Li H, Sun H. Coordination chemistry of metals in medicine: Target sites for bismuth. Coord Chem Rev. 1999;185–186: 689–709. doi: 10.1016/s0010-8545(99)00018-1

60. Chen Z, Zhou Q, Ge R. Inhibition of fumarase by bismuth(III): Implications for the tricarboxylic acid cycle as a potential target of bismuth drugs in Helicobacter pylori. BioMetals. 2012;25: 95–102. doi: 10.1007/s10534-011-9485-7

61. Beatrix Bialek FT, Bialek B, Hensel R. Medical Use of Bismuth: the Two Sides of the Coin. J Clin Toxicol. 2011;s3: 1–5. doi: 10.4172/2161-0495.S3-004

