## Supplementary materials for "Bismuth nanoparticles obtained by a facile synthesis method exhibit antimicrobial activity against *Staphylococcus aureus* and *Candida albicans*"


Roberto Vazquez-Munoz [1,a], M Josefina Arellano Jimenez[2], Jose Lopez-Ribot[1]

[1] The University of Texas at San Antonio, San Antonio, TX, USA

[2] The University of Texas at Dallas, San Antonio, TX, USA

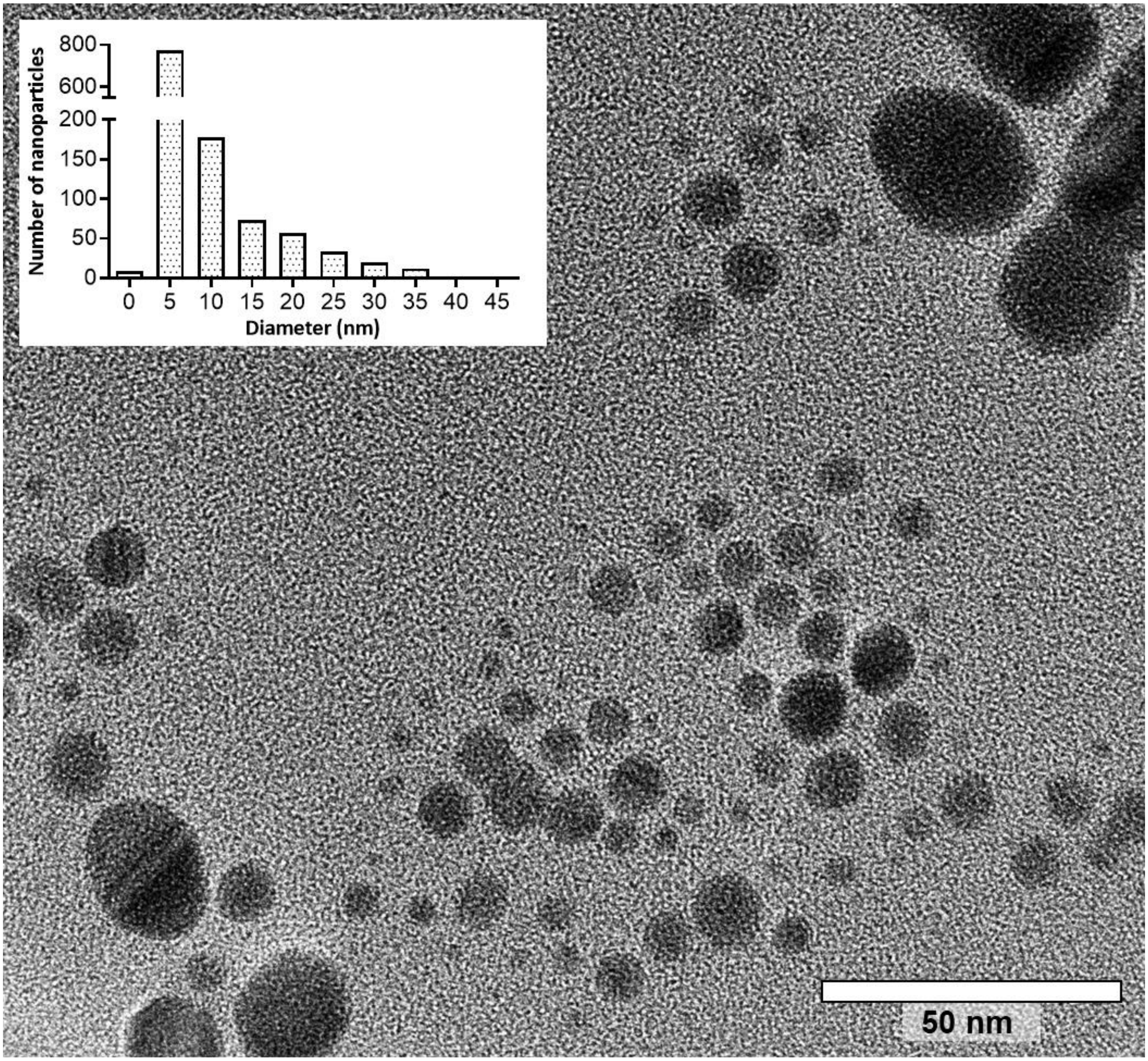


**Fig. S1. TEM images confirm the presence of BiNPs**. PVP-BiNPs display an aspect ratio close to 1 and the size is below 10 nm. Occasional other shapes and sizes are also observed.

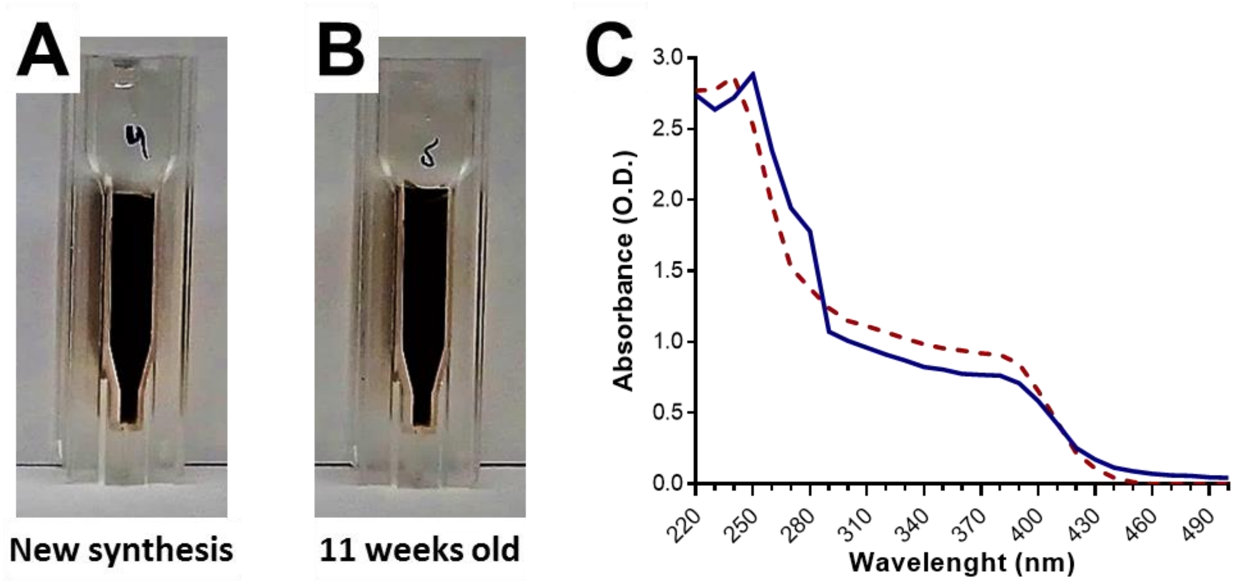


**Fig. S2. Bismuth nanoparticles in aqueous solutions remain stable over time**. Visual examination reveals that BiNPs remained stable over time. Visually, an 11-week-old synthesis (B) kept light-protected at 4 $^{o}$C, appears identical to a freshly prepared synthesis (A). UV-Vis spectrophotometry reveals that their optical properties (absorbance profiles) remain similar -with minor changes- (C) between the old synthesis (red) and the new preparation (blue).

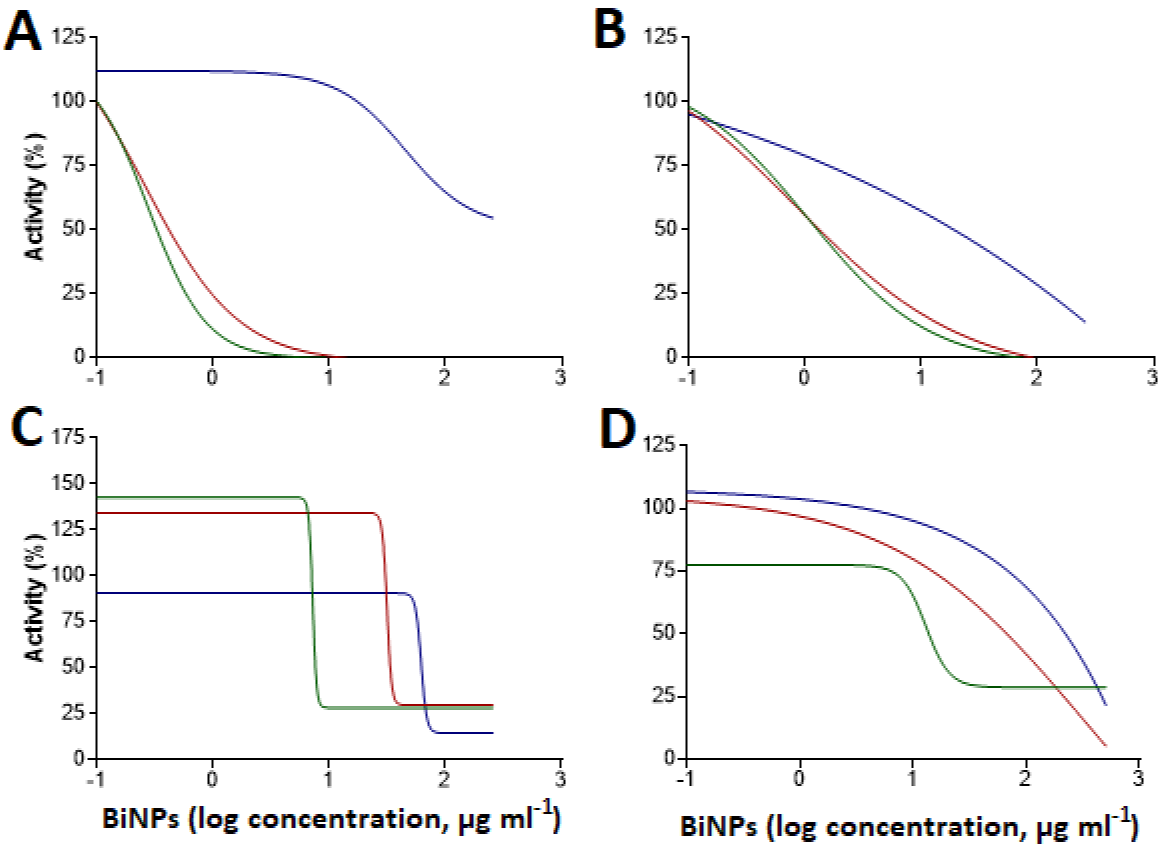


**Fig. S3. Inhibition of *S. aureus* and *C. albicans* by different bismuth compounds under both planktonic and biofilm growing conditions.** The dose-response curves against *S. aureus* (top panels), and *C. albicans* (bottom panels) show that bismuth compounds display different degrees of antimicrobial activity under planktonic (A, C) and biofilm (B, D) growing conditions. Lines: green (BiNPs), red (bismuth-BAL), and blue ($Bi(NO_3)_3$).
